# Genome size and ploidy of the German species of *Veronica* L. (*Plantaginaceae*)

**DOI:** 10.1101/2023.12.22.573074

**Authors:** Dirk C. Albach, Mareike Daubert

## Abstract

Chromosome numbers and genome size estimates provide essential information for the differentiation of plant species. Especially, closely related species that are morphologically difficult to distinguish are often easily distinguishable by genome size. Flow cytometry has facilitated in recent years the detection of such differentiation at the genomic level. It further helped understand the distribution of ploidy levels within species. The genus *Veronica* includes 37 species in Germany including some taxonomically challenging species groups and some species with intraspecific variation in ploidy. We, here, present 36 new genome size estimates and 44 estimates of ploidy, six and seven, respectively, from Germany, for these 37 species. Estimates of *V. aphylla, V. alpina, V. fruticans*, and *V. fruticulosa* are first estimates for the species. These estimates provide an important basis for future studies on the genus. Some examples are discussed in more details, such as the distribution of ploidy levels in *V. longifolia* and *V. chamaedrys* in Germany or the importance to study ploidy levels in *V. satureiifolia* and relatives in southwestern Germany.

---

Chromosomes were first described by Flemming in 1882 (O’Connor & Miro 2008). It was soon found that the number of chromosomes per nucleus is mostly a stable character within species and often variable among species, which is useful for taxonomic purposes (Stuessy 2009). Consequently, scientists have accumulated more than 70,000 counts for all kind of angiosperms, covering about 20% of all species (Rice eta l. 2015). This data has been used not just for taxonomic purposes but also to infer ancestral chromosome numbers (20962), models for the mechanistic evolution of chromosomes (21360), and ecological correlates (15134). However, there is still a dearth of understanding the evolutionary forces selecting for higher or lower chromosome numbers. While it is easily conceivable that organisms with a large number of chromosomes have problems during meiosis, some species have a remarkably large number of chromosomes (Rice & al. 2015). Larger numbers and large variation among taxa are mostly considered to affect gene flow and produce reproductive barriers (Bowers and Patterson 2021). This provides some theoretical support for the use of chromosome number as a taxonomic character.

Whereas basic chromosome numbers are considered mostly uniform for a species and even species groups, individuals and closely related species may still vary in chromosome numbers based on changes in ploidy level. Polyploidy, caused by whole genome duplications, occur either as a result of intraspecific crossing, potentially even in a selfing individual, or interspecific hybridization, potentially even among different genera (Stuessy 2009). Polyploidy, similar to other chromosome number changes, confers at least some reproductive isolation (Baack & al. 2015). Ploidy levels can be inferred by chromosome number counts or, more commonly, in recent years, by flow cytometry, increasing enormously our understanding for the distribution of intraspecific ploidy variation (e.g., Suda & al. 2007; Bardy & al. 2010).

Flow cytometry provides information beyond the ploidy level, which is exact genome sizes. Apart from whole genome duplications, genome size may be influenced by small-scale duplications or deletions. The largest effect of these have activation or inactivation and deletion of transposable elements (Petrov 2002). Such processes occur continuously and lead to minor differences in genome size among individuals. However, larger variation (more than 5%) is rare. Intraspecific variation in genome size is a hotly debated topic. Establishing genuine variation is difficult due to technical variation and variation caused by secondary metabolites (Greilhuber 1998). However, given the selective effect of genome size related to environmental effects, nutrient availability and growth strategies (Bennett 1972; Hessen & al. 2008; Müller & al. 2019), it is more than likely that adaptive variation in genome size exists within species (e.g., Bilinski & al. 2018). Thus, genome size and its intraspecific variation is important karyological information that may help understand evolutionary forces shaping genomic features but also aid in taxonomic conclusions (e.g., Prancl & al. 2015).

The genus *Veronica* is the largest genus of Plantaginaceae with about 450 species, approximately 180 species occurring exclusively in New Zealand where they form the largest plant radiation. In Germany, the genus is represented by 37 species (Albach 2021). Some of these species are difficult to distinguish and have caused heated taxonomic debates in the past. Since the first DNA-based phylogenetic analysis focusing on relationships among Northern Hemisphere species of *Veronica* (Albach and Chase, 2001), these debates have received new momentum. Several of these species have been analyzed in more detail now with various molecular methods and flow cytometry, so that most debates have been settled and we now have an overview on the German species of the genus and which taxonomic questions are still open.

We, here, summarize our current knowledge on the phylogenetic relationships of the German species, provide a review of ploidy levels including several new estimates and, finally, highlight open taxonomic questions.

## Materials and Methods

Nomenclature for the genus *Veronica* follows Müller & al. (2021). Information on chromosome numbers were investigated in the literature. The literature up to 2007 was summarized by Albach & al. (2008). Therefore, information up to that time was taken from that publication and more recent counts were searched from literature and online databases (e.g. Rice & al. 2015; Paule & al. 2017).

Ploidy levels and genome sizes were estimated following standard protocols established in our lab (Meudt & al. 2015). In particular, we used mainly *Solanum pseudocapsicum* L. (1C =1.2946pg; Temsch & al. 2010) but sometimes also *Hedychium gardnerianum* Sheph. ex Ker Gawl. (1C = 2.01pg) or Zea mays L. ’CE-777’ (1C=2.7165; Lysák & Dolzel 1998) as internal standards. We used the OTTO buffer and stained at 4°C for 1 hour with propidium iodide. Measurements were conducted on a Partec CyFlow SL (Münster, Germany). Measurements per sample included 5000 particles and at least 500 particles per counted peak.

## Results & Discussion

We found 90 chromosome counts based on German plants (Suppl. File 1). Five are not included in the database of Paule & al. (2017). These are not included because their origin from German plants is not sufficiently clear. The two counts of *V. hederifolia* by Hofelich (1935) are from plants from the Botanical Garden in Tübingen but it is not clear whether they are spontaneous (as we believe) or cultivated. The counts by Tischler (1937) for *V. persica* and *V. polita* are not documented clearly but likely derive from plants from Northern Germany. The origin of the seeds used for the count of *V. opaca* by Beatus (1936) are either from Germany or Czech Republic. These counts represent 22 of the 37 species. However, for all species counts exist from somewhere in the range.

We here provide 36 new genome size measurements plus an additional 44 estimates of ploidy (unrepeated genome size estimates or estimates with cv above 7) using flow cytometry. Of these, six genome size (of six species) and seven ploidy estimates (of four species) are from German plants. The estimates of three species (*V. aphylla, V. fruticans, V. fruticulosa*) are first genome size estimates of the species. With the values published here, genome size estimates are available for all species of *Veronica* in Germany except *V. anagalloides* and *V. opaca*. However, only for ten of 37 species estimates are from German plants. Nevertheless, the study provides the basis for easy determination of ploidy by flow cytometry for all German species or species groups with variation in ploidy.

### 1. Veronica montana

*Veronica montana* is a widespread European species but no chromosome counts and no genome size estimates are available for German plants. However, all counts, many from neighboring countries (Netherlands, France, Czech Republic, Poland) indicate it is a diploid (2n = 18) species (Albach & al. 2008). This is also confirmed by genome size estimates from Britain and the Netherlands (Bennett & Smith 1991; Zonneveld 2019). However, those two estimates differ by 15% (1C = 0.73 vs. 0.85 pg), necessitating further analyses to establish firmly the genome size of this species. Since both estimates are intermediate between those of the phylogenetically related species (Meudt & al. 2015), there is currently no suggestion about the more correct value.

### 2. Veronica scutellata

*Veronica scutellata* is universally diploid (2n = 18; Albach & al. 2008) and so is the only German count (Scheerer 1939). Also, our genome size estimates confirm this ploidy level, although they are higher than in other species of the subgenus. Despite continuity in ploidy level, there may be some intraspecific variation in genome size. Our Ukrainian estimate (Table 1) does not differ from the one from the Netherlands (1C = 0.885pg; Zonneveld 2019) but both differ from our Siberian plants by about 5%, although the latter is not reliable.

**Table 1:** Information on genome size measurements for *V*. subgen. *Veronica*. Genome size measurements with only one run and/or cv above 7 were intended as tests of ploidy and should therefore not be considered reliable genome size estimates.

| Species | Internal standard | CV/runs | Genome size (1C in pg) | Ploidy | Voucher | Locality |
| --- | --- | --- | --- | --- | --- | --- |
| *Veronica scutellata* | *Solanum pseudocapsicum* | 5.90-6.38/3 | 0.939 | 2x | Albach & Kosachev 1321, OLD | Russia, Altaisky Kray, Barnaul, near Bot. Gard. |
|  | *Solanum pseudocapsicum* | 7.83/1 | (0.883) | 2x | Albach & al. 1458, OLD | Ukraine, Kyiv, Desna river, near Prinovo |
| *Veronica aphylla* | *Solanum pseudocapsicum* | 3.16-3.19/3 | 0.483 | 2x | *Albach S524*,<br>OLD | Austria,<br>Steiermark,<br>Hochschwab<br>gruppe,<br>Griesmauer;<br>ex BG<br>München |
| *Veronica urticifolia* | *Solanum pseudocapsicum* | 2.44-4.74/3 | 0.633 | 2x | *Albach S1039*,<br>OLD | Switzerland,<br>Chambesy;<br>ex BG<br>Geneve |
| | *Solanum pseudocapsicum* | N.N.($<5$)/3 | 0.633 | 2x | *Albach S1064*,<br>OLD | Austria,<br>Carinthia,<br>North slope<br>of Sechter;<br>ex BG<br>Klagenfurt |
| *Veronica alpina* | *Solanum pseudocapsicum* | 2.53-3.75/3 | 0.691 | 2x | *Albach S1005*,<br>OLD | Iceland,<br>Lystigardur<br>Akureyrar; ex<br>BG Akureyri |
|  | *Solanum pseudocapsicum* | 3.72-4.20/3 | 0.714 | 2x | *Albach S689*,<br>OLD | Iceland,<br>Landsmanna<br>afrettur; ex<br>BG Münster |
| *Veronica bellidioides* | *Solanum pseudocapsicum* | 2.08-2.15/3 | 1.070 | 4x | *Albach S507*,<br>OLD | France,<br>Galibier; ex<br>BG Grenoble |
|  | *Solanum pseudocapsicum* | 2.14-3.14/3 | 1.085 | 4x | *Albach S1004*,<br>OLD | Cultivated;<br>ex<br>Gesellschaft<br>der<br>Staudenfreu<br>nde |

### 3. Veronica officinalis

The type species of the genus, *V. officinalis* is widespread across Eurasia. Böcher (1944) counted two German plants as tetraploid, which agrees with most other of the close to 100 counts (Albach & al. 2008). Diploids are found in Portugal, Gotland (Sweden), and Siberian Sayan Mountains and Irkutsk (Albach & al. 2008).

Previous genome size measurements range between 1C=1.01-1.10pg (Siljak-Yakovlev et al. 2010; Pustahija et al. 2013; Castro et al. 2012; Bai et al. 2012). This range is mostly confirmed with just one reliable value a bit lower (Table 2). In preparation of a wider phylogeographical analysis of the species, we sampled more broadly across Europe but only found tetraploids (Table 2). Based on the diploid count from Gotland we anticipated a wider diploid lineage in the Baltic area. However, from Poland and Lithuania only tetraploid populations are known so far. This is also confirmed by four further measurements in which a clear separation of standard and sample peaks was not possible due to their close proximity (results not shown). There are two ploidy estimates from Germany, one from the Solling mountains and one from Saxony, which are tetraploid.

**Table 2:**
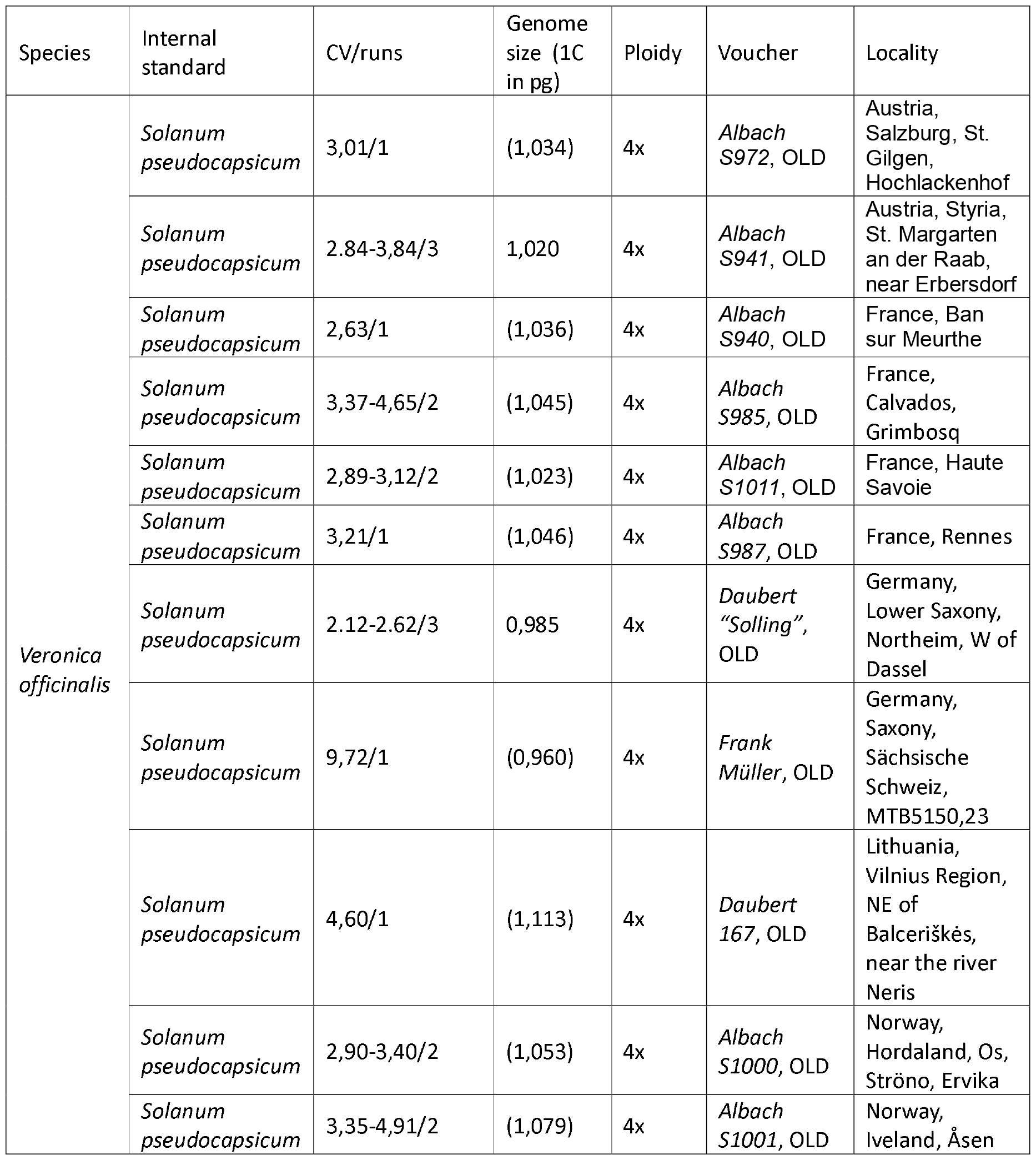
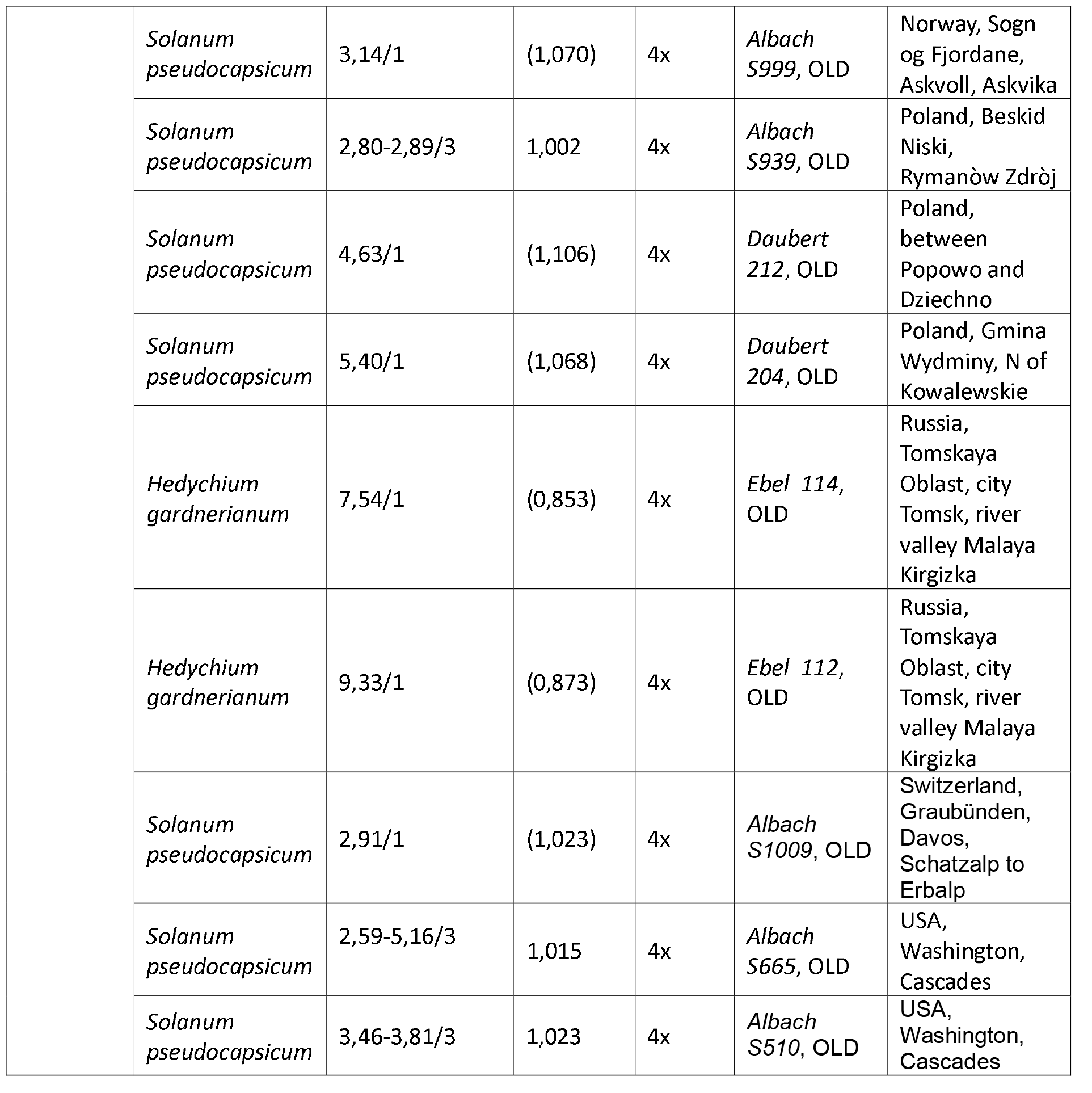
Information on genome size measurements for *V. officinalis*. Genome size measurements with only one run and/or cv above 7 were intended as tests of ploidy and should therefore not be considered reliable genome size estimates.

### 4. Veronica aphylla

No chromosome count is available for this species from Germany, where it occurs marginally in the Alps. All eleven counts from the Alps and Carpathians indicate that it is a diploid species (2n = 18; Albach & al. 2008). Our genome size estimate from Austria (1C=0.483pg) is the first for the species and comparable to that of the closely related *V. baumgartenii* (Albach & Greilhuber 2004).

### 5. Veronica urticifolia

Similar to *V. aphylla*, no chromosome count is available for German plants of this montane species but all other nine counts from throughout Europe are diploid (2n = 18; Albach & al. 2008). Genome size estimates for the species are likewise not from Germany. The first estimate (1C = 0.643 pg; Albach and Greilhuber 2004) was from a plant of unknown origin but our new estimates are in the same range (Table 1). However, these are about double those from Pustahija & al. (2013) and Siljak-Yakovlev (2010), although Pustahija & al. (2013) also reported estimates in our range, which they interpreted as tetraploid. It, therefore, remains an open topic whether diploids and tetraploids exist next to each other in this species or whether the lower or higher numbers are artefacts. The latter hypothesis is supported by the fact that the genome size per chromosome set (1Cx-value) would be 35-50% lower than in related species (e.g., *V. alpina*; Albach & al. 2007).

### 6. Veronica alpina

*Veronica alpina* is another alpine species without a chromosome count from Germany but with 38 counts from throughout the range supporting its diploid status (2n=18) and only a single tetraploid count (Albach & al. 2008). Its genome size (1C=0.691-0.714pg; Table 1) has previously not been reported but is only slightly smaller than the one reported for the closely related *V. copelandii* (Albach & Greilhuber 2004).

### 7. Veronica bellidioides

The next alpine species without a chromosome number is *V. bellidioides*. In contrast to the previous species, more than 40 chromosome numbers from throughout the range suggest that it is a tetraploid species with diploids occurring in the Pyrenees (2n=36; Albach & al. 2008). Our new genome size estimates (Table x) are slightly higher (3-4%) than those reported earlier (Albach & Greilhuber 2004), which is to be expected based on the different methods used (Feulgen densitometry versus flow cytometry) as discussed by Meudt & al. (2015).

### 8. Veronica acinifolia

*Veronica acinifolia* is a diploid (2n=14), annual species, which has become rare in Central Europe due to changes in agricultural practices. The count from Germany (Hand & Gregor 2015) is from Baden-Württemberg and agrees with all other 12 counts from elsewhere (Albach & al. 2008). The three accessions from which genome size was measured all originated from Nantes in France (Table 3). Of these, two are very similar but the third has about 10% less DNA. The published estimate (Castro & al. 2012) is intermediate between these two. So, further investigation is necessary to find the exact size, although no change in ploidy is expected.

**Table 3:**
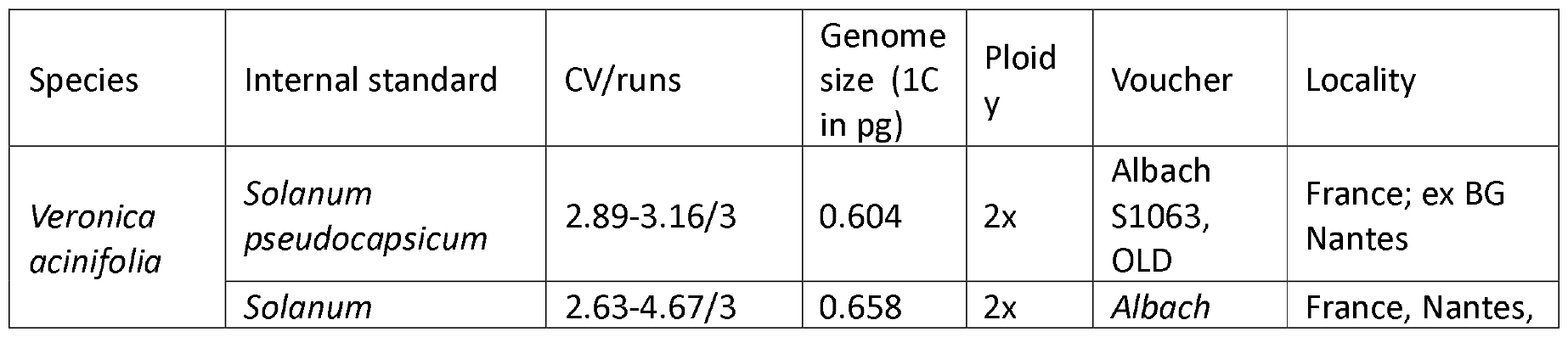
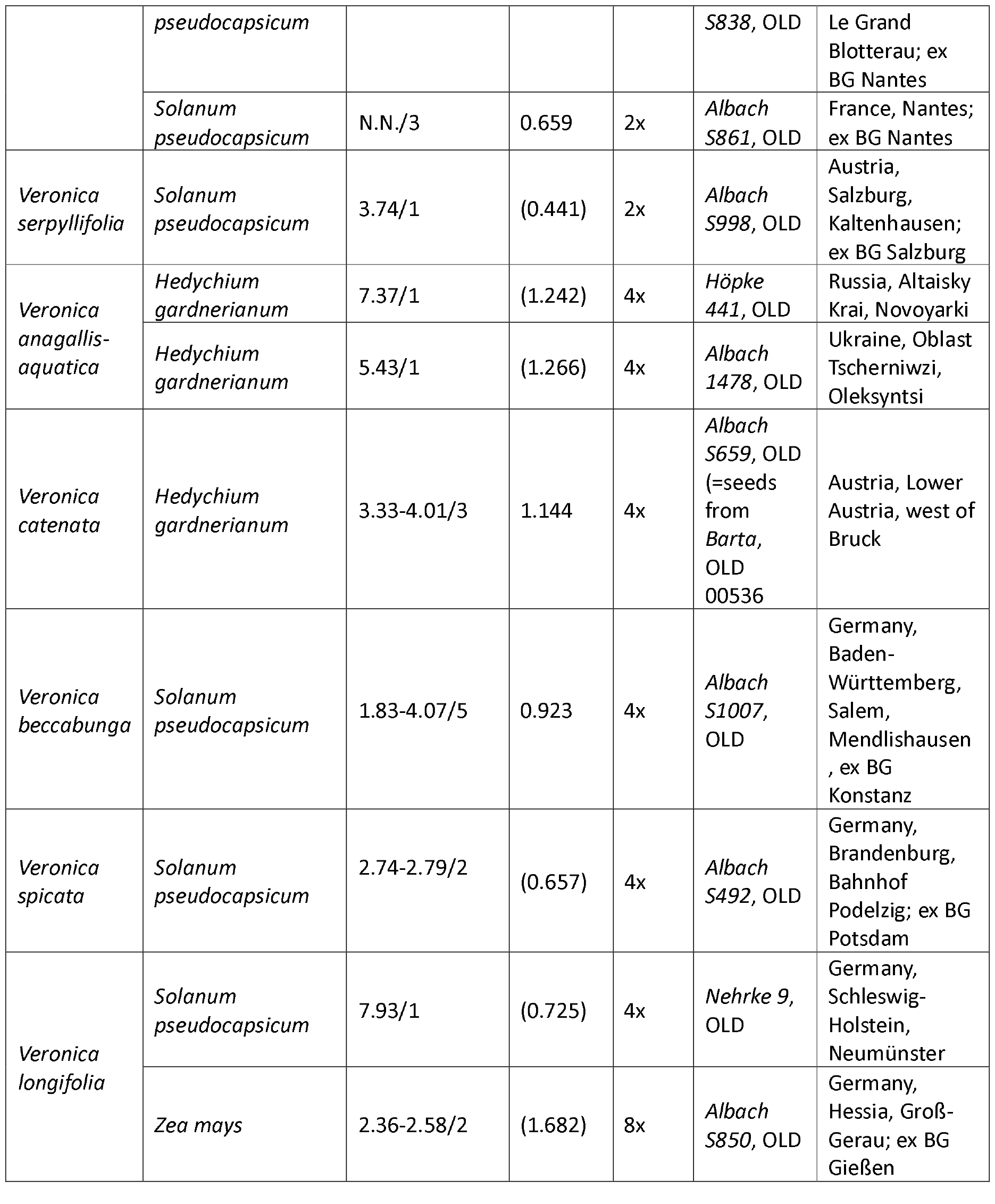
Information on genome size measurements for *V*. subgen. *Beccabunga* and *Pseudolysimachium*. Genome size measurements with only one run and/or cv above 7 were intended as tests of ploidy and should be therefore not be considered reliable genome size estimates.

### 9. Veronica serpyllifolia

There are more than 70 chromosome counts for this species with all except two being diploid including both German counts (2n =14; Albach & al. 2008). Genome size of about 1C=0.44-0.45pg was first estimated based on a plant from Georgia based on Feulgen densitometry (Albach & Greilhuber 2004) and later confirmed by flow cytometry (Castro & al. 2012). Our estimate here from Austria agrees with these earlier estimates (Table 3). So, it can be expected that all German populations are diploid with approximately that genome size.

### 10. Veronica peregrina

*Veronica peregrina* is hexaploid but as discussed in Albach & al. (2008) two chromosomes may have been fused to reduce the number of chromosomes to 2n =52. This is the number reported from many countries (but not Germany) in about 16 publications summarized in Albach & al. (2008). The first genome size estimate of the species (1C=0.95pg) was published by Albach & Greilhuber (2004) based on plants growing spontaneously in the Botanical Garden Frankfurt. The subsequent publications from Portugal and Netherlands were either a bit larger (1C=0.98pg; Castro & al. (2012); 1C=1.03pg; Zonneveld 2019) or the one from Germany (Oldenburg) smaller (1C=0.87pg; Meudt & al. 2015). The variation of almost 20% may be of technical nature or related to the high ploidy and variation in genome downsizing. The nature of genome size variation and possibly chromosome number variation should be further investigated.

### 11. Veronica anagallis-aquatica

*Veronica anagallis-aquatica* is the most widespread species of the aquatic species in the genus with an almost cosmopolitan distribution now. Despite the wide distribution area and chromosome counts from many parts of the range, the species is tetraploid (2n = 36; Albach & al. 2008) with the exception of some northern Indian counts (Kaur & Singhal in Marhold 2012). The three German counts (Schlenker 1936; Paule & al. 2017) agree with this. The first genome size estimate (Albach & Greilhuber 2004) was 1C = 1.08 pg with a second publication (Zonneveld 2019) resulting in a 23% larger estimate. Our estimates here (Table 3) indicate that the first estimate may, indeed, have been somewhat small.

### 12. Veronica catenata

The morphologically similar *V. catenata* is largely sympatric with *V. anagallis-aquatica* in Europe and North America but does not reach further into Asia and its occurrence in North Africa is dubious. It is also tetraploid (2n = 36) throughout its range (Albach & al. 2008) and so is its only count from Germany (Dersch in Paule & al. 2017). Zonneveld (2019) provided the first genome size estimate (1C = 1.235 pg) with our estimate (Table 3) being a bit lower. It remains to be tested further whether *V. catenata* has a consistently smaller genome than *V. anagallis-aquatica*. It would be very helpful to distinguish these two species, which are very difficult to differentiate, especially in vegetative state.

### 13. Veronica anagalloides

The aquatic species of *Veronica* are notoriously difficult to determine due to large plasticity related to water availability (Ellmouni et al 2017). Species limits in Egypt and Near East are not understood, yet (Ellmouni & al. 2018; Hosseinnejad Azad & al. 2020). Thus, chromosome counts from these areas are sometimes difficult to assign to a species. Of the 37 counts available (Albach & al. 2008; Sanchez Agudo & al. 2011; Probatova & al. 2014) 25 are diploid (2n = 18). Most of the tetraploid counts are from Asia and may refer to a different taxon than the Mediterranean taxon. Whereas misidentification cannot be ruled out, especially the tetraploid counts from Spain (Sanchez-Agudo & al. 2014) suggest that there may be another tetraploid, strongly glandular lineage in *V. anagalloides*. In central Europe, counts (all diploid) are available from Austria, Hungary, Slovakia, and France (Marchant 1970, Peniastekova in Majovsky & al. 2000, Schlenker 1936). So, this is the expected chromosome number of the German plants. No genome size estimate is available for the plant, but it is expected to be about half of that of *V*. anagallis-aquatica (1C = ca. 0.6pg).

### 14. Veronica beccabunga

No chromosome count of *V. beccabunga* from Germany is available despite more than 60 throughout the range (Albach & al. 2008). Most of the counts are diploid (2n=18) but tetraploid counts (2n=36) have been reported from Italy, Poland, Spain, and Sweden. The closely related species *V*. americana is likewise tetraploid and the exact relationship between both species is not known. Therefore, more information on the distribution of cytotypes would be desirable. This would be facilitated by a reliable genome size for flow cytometrical determination of ploidy. However, published genome sizes vary by around 50%. Bennett and Smith (1991) reported a genome size of 1C=0.73 pg. Subsequent estimates have been larger between 1C=0.805pg (Zonneveld 2019) to 1C=1.23 pg (Hidalgo & al. 2015). Our estimate here (Table 3) is intermediate with 1C=0.923pg. Therefore, it would be important to have chromosome counts of individuals for which genome size is measured in parallel. Comparison with related species does not help in the case since no genome size estimate for *V. americana* is available and those of *V. anagallis-aquatica* and *V. catenata* (see above) are either lower (diploid; 1C=0.5-0.6pg) or higher (tetraploid; 1C=1.08-1.33pg) with only the estimate of Hidalgo & al. (2015) in the range of tetraploid *V. anagallis-aquatica*. This suggests that this estimate is for tetraploid plant supporting the presence of tetraploids based on chromosome counts (Sanchez Agudo & al. in Marhold & Breitwieser 2011).

### 15. Veronica spicata

Two chromosome counts for German *V. spicata* are available showing the octoploid level (Huber 1927; Dersch in Paule & al. 2017). Note that the second reference provides the unusual number 2n=64, whereas all other publications for this species provide 2n=68 (Albach & al. 2008). So, it is likely to be miscounted. Both reports are from southwestern Germany and so is the published genome size measurement from Brey (Rhineland-Palatia; Buono & al. 2021). In contrast, our report here (Table 3) is tetraploid and from eastern Germany. This fits with the continent-wide analysis of Buono & al. (2021), who reported a tetraploid plant from Poland. Genetically, the octoploid West German *V. spicata* belong to the tetra- and octoploid Western European lineage of *V. spicata*. The closest tetraploids of that lineage occur in the Alsace. In contrast, the tetraploid Polish and likely the East German *V. spicata* belong to the Eurasian lineage (Buono & al. 2021). The two lineages apparently border in Germany, and it will take more intensive analysis about their exact distribution pattern in Germany. The situation is, however, more complicated by the presence of a tetraploid Pannonian-Balkan lineage, which extends to Austria and the Czech Republic and possibly into Bavaria (Buono & al. 2021). The Baltic octoploids have not been analyzed genetically and may even constitute a fourth genetic lineage reaching Germany.

### 16. Veronica longifolia

Despite being widespread in Germany, there is no chromosome count for this species available from Germany. Our measurements (Table 3) are the first information on ploidy for the species from Germany. Generally, the octoploid level predominates in the species with tetraploids occurring scattered in three of the four lineages detected by Buono & al. (2021), although they are not genetically as clearly differentiated as in *V. spicata* (Buono & al., 2021). Tetraploids occur in the Pannonian lineage in Hungary, in the eastern European-Siberian lineage tetraploid specimens from European Russia, eastern Anatolia and Siberia, and in the Baltic lineage but they have not been detected in the Western European lineage. Our tetraploid sample from Schleswig-Holstein may consequently belong to the Baltic lineage, whereas the octoploid Hessian is more likely to belong to the Western European lineage. However, this conclusion should be considered carefully since these lineages are not geographically completely coherent, likely due to horticultural spread of the species, as indicated by the presence of specimen from the Pannonian lineage in the outskirts of Mannheim (Buono & al. 2021).

### 17. Veronica fruticans

*Veronica fruticans* is widespread in the alpine regions of Europe and is uniformly diploid (2n = 16; Albach & al. 2008). It reaches Germany in the Alps and the Black Forest but no count from that area is available. Our genome size estimate (Table 4) is the first for the species and is only slightly higher than that of related species from the Balkan Peninsula (Albach & al. 2009).

**Table 4:** Information on genome size measurements for *V*. subgen. *Stenocarpon, Pellidosperma* and *Chamaedrys* (without *V. chamaedrys*).

| Species | Internal standard | CV/runs | Genome size (1C in pg) | Ploidy | Voucher | Locality |
| --- | --- | --- | --- | --- | --- | --- |
| *Veronica fruticans* | *Solanum pseudocapsicum* | 2.76-3.12/3 | 0.938 | 2x | Albach S710, OLD | Switzerland, Glarus, Bergstock; ex BG München |
| *Veronica fruticulosa* | *Solanum pseudocapsicum* | 2.92-3.45/3 | 1.000 | 2x | Albach S1003, OLD | Cult., Gesellschaft der Staudenfreunde |
| *Veronica triphyllos* | *Solanum pseudocapsicum* | 2.47-2.67/3 | 0.587 | 2x | Albach S1049, OLD | Cult. ex BG Wien |
|  | *Solanum pseudocapsicum* | 2.36-2.73/3 | 0.588 | 2x | Albach S434, OLD | Russia, Altay, ex BG Osnabrück |
| *Veronica praecox* | *Solanum pseudocapsicum* | 3.75-4.13/3 | 0.541 | 2x | Albach S852, OLD | France, Prelles; ex BG Grenoble |
|  | *Solanum pseudocapsicum* | 3.30-6.24/3 | 0.542 | 2x | Albach S1046, OLD | France, Prelles; ex BG Grenoble |
| *Veronica arvensis* | *Solanum pseudocapsicum* | 2.90-4.50/3 | 0.423 | 2x | Albach S858, OLD | France, Lautaret; ex BG Grenoble |
|  | *Solanum pseudocapsicum* | 4.16-5.86/3 | 0.436 | 2x | Albach S859, OLD | Germany, Northrhine-Westphalia, Lengerich station; ex BG Münster |
| *Veronica dillenii* | *Solanum pseudocapsicum* | 2.99-3.53/3 | 0.612 | 2x | Albach S682, OLD | Austria; ex BG Wien |
|  | *Solanum pseudocapsicum* | 3.04-4.7/6 | 0.632 | 2x | Albach S839, OLD | Austria, Zogelsdorf |

### 18. Veronica fruticulosa

Similar to the case of *V. fruticans*, there is no chromosome count available from Germany for this species from the Alps, which is otherwise uniformly diploid (2n = 16; Albach & al. 2008). Again, our genome size estimate is the first for the species (Table 4) and it is slightly higher than that of *V. fruticans* and even higher than that of related species (Albach & al. 2009).

### 19. Veronica triloba

*Veronica triloba* has long been recognized as distinct from *V. hederifolia* despite being closely related (Opiz 1825). The morphological similarity has led to considerable undersampling and uncertainty in its distribution in Germany, but it is certainly more restricted to the southern half of Germany (Albach 2021). Fischer (1967) helped in the differentiation of the taxon and established that it is a diploid taxon. Chromosome counts and genome size measurements from Germany are still lacking but the genome size of Turkish plants has been published by Albach & Greilhuber (2004).

### 20. Veronica sublobata

In the same publication, in which Fischer (1967) established the diploid nature of *V. triloba*, he also raised *V. hederifolia* subsp. *lucorum* to the species level, as *V. sublobata*, establishing it as a tetraploid relative of the hexaploid *V. hederifolia*. No chromosome count from Germany is available, but Albach & al. (2008) summarized 60 counts from throughout the rest of the range supporting the results of Fischer (1967). The first genome size measurement for one of the two species was by Albach & Greilhuber (2004), who reported a 1C-value of 1.41. Based on comparison with the related *V. cymbalaria* they assumed that it was a hexaploid plant. Subsequently, Castro & al. (2012) reported a 1C-value of 2.08 from Portuguese plants of *V. hederifolia*, which suggested that the original estimate was a tetraploid plant. This view has been confirmed by close inspection of the voucher specimens (M.A. Fischer, pers. Comm.). Zonneveld (2019) reported 1C-values of 1.55pg and 2.06pg for subsp. *hederifolia* and subsp. *lucorum*, respectively, which suggests that *V. sublobata* is hexa- and *V. hederifolia* is tetraploid. Based on the available chromosome numbers, many more flowcytometrical measurements and phylogenetic evidence of DNA data (Albach & Kur, in prep.), we believe that Zonneveld (2019) turned his values around and that *V. sublobata* is a European tetraploid lineage. Its distinctiveness in Central Europe is well-established but the southern and eastern borders of its distribution area are unclear (Fischer 1975, Albach & Kur, in prep.).

### 21. Veronica hederifolia

*Veronica hederifolia* is hexaploid (2n=54) in Germany based on seven counts (Albach & al. 2008; Buttler in Paule & al. 2017). As discussed above under *V. sublobata* its genome size (1C-value) is between 2.00-2.10(-2.20) pg (Castro & al. 2012; Albach & Kur, in prep.). Apart from its karyological difference, in Germany it can be morphologically differentiated from its close relatives (Fischer 1967, 1975; Atha & al. 2021; Albach 2021; Albach & Kur, in prep.) and is a genetically separate lineage (Albach & Kur, in prep.). The questions around its circumscription should focus on southern and eastern Europe, as well as Turkey to Iran.

### 22. Veronica cymbalaria

Both, tetraploid and hexaploid populations are known from this species (Albach & al. 2008). However, only tetraploids are known from Germany (2n = 36; Hügin & Hügin 2002). Albach (2007) has demonstrated that in the Mediterranean tetraploids have formed at least twice from the close relatives *V. panormitana* and *V. trichadena* and hexaploids have formed multiple times from backcrossing of the tetraploids with either parent. Future investigations may discover hexaploids from Germany as well, which are widespread in the Mediterranean as well (Albach 2007). Their expected genome sizes (1C-value) are 0.4pg, 0.8pg and 1.4-1.5pg for the di-, tetra- and hexaploid level, respectively.

### 23. Veronica triphyllos

*Veronica triphyllos* is a diploid (2n = 18) annual species, which is widely distributed but not common in Germany. The chromosome number is based on 18 counts throughout the range including one from Germany (Hofelich 1935). In contrast to its chromosome number its genome size is less clear. Albach & Greilhuber (2004) reported 1C=0.71pg for Turkish plants. Later, Zonneveld (2019) reported 1C=0.485pg for Dutch plants. Our estimates here (Table 4) are intermediate between these (1C = 0.59pg). More measurements are necessary to see whether there is a technical artefact in the Feulgen-based estimate of Albach & Greilhuber (2004; Meudt & al. 2015) or whether there is geographic variation.

### 24. Veronica praecox

No chromosome count from German plants is available for *V. praecox*, a warmth-loving annual species, which is diploid (2n = 18) according to 19 chromosome counts throughout its range (Albach & al. 2008). Based on two measurements, its genome size is between 1C=0.54pg (Table 4) and 1C=0.59pg (Zonneveld 2019).

### 25. Veronica arvensis

*Veronica arvensis* is one of the most widespread weedy species of the genus and its chromosomes have been counted 48-times (Albach & al. 2008; Doostmohammadi & al. 2021; Sanchez-Agudo & al. in Marhold & Breitwieser 2011; An’kova & al. In Marhold & Kucera 2019) including two counts from Germany and all counts are diploid (2n=ca. 16). Also, its genome size has been measured multiple times. The earliest estimate by Albach & Greilhuber (2004) was too low, likely due to problems with the Feulgen densitometry (Meudt & al. 2015) but all subsequent analyses, including our estimate from Germany (Table 4), support a genome size of about 1C= 0.41-0.45pg (Meudt & al. 2015, Castro & al. 2012, Zonneveld 2019, Table xxx).

### 26. Veronica verna

*Veronica verna* is an annual species, closely related with *V. arvensis*. It is likewise diploid (2n = 16) based on 21 counts (Albach & al. 2008; Sanchez-Agudo & al. in Marhold & Breitwieser 2011; Doostmohammadi & al. 2021) including one from Germany (Hofelich 1935). Albach & Greilhuber (2004) reported a genome size of 1C= 0.54pg for a plant from Georgia using Feulgen densitometry. Due to the problems of the method, further estimates are necessary (Meudt & al. 2015), especially to confirm the difference in genome size to the closely related *V. dillenii* (see below).

### 27. Veronica dillenii

Chromosome number data is similar for *V. verna* as for the closely related *V. dillenii*. Both are diploid (2n = 16) based in the latter case on 13 counts including one from Germany (Albach & al. 2008; Sanchez-Agudo & al. in Marhold & Breitwieser 2011). Both species are vegetatively very similar but differ in their flower size and possibly in their phytochemical arsenal (Albach 2021). Given their close relationship it is rather surprising that their genome size differs by 15% (1C = 0.61-0.63pg vs. 0.54pg; Table 4, Albach & Greilhuber 2004). Apart from technical reasons or geographic variation, there may be a biological reason. Albach & Greilhuber (2004) reported that selfing species of *Veronica* have a significantly smaller genome than outcrossing species. Given the close correspondence of flower size with outcrossing rate in *Veronica* (Scalone & al. 2013), our hypothesis is that the larger flower and higher outcrossing rate of *V. dillenii* is the reason for a larger genome. In line with that hypothesis, the strongly selfing *V. arvensis* has a lower genome size and the obligate outcrosser *V. chamaedrys* has a larger genome size.

### 28. Veronica chamaedrys

Possibly the best investigated species of the genus, there are about 180 counts available for the species with most of them tetraploid (2n = 32; Albach & al. 2008). Diploids are found scattered among the tetraploids on the large continental scale and sometimes even mixed in one population (Fischer & Mirek 1986; Bardy & al. 2010). Diploids are predominantly in southeastern Europe reaching north into Byelorussia (Dzhus & Dmitrieva 2001), southern Poland (Fischer & Mirek 1986) and southern Germany (Lippert & Heubl 1989). So far, there are 14 chromosome counts from Germany, eight of them diploid and all of these from the southern half of Bavaria. Based on AFLP and flow cytometry, Bardy & al. (2010) detected three lineages on the Balkan Peninsula (west, east, south), all of which contained diploids and tetraploids, which originated independently in all three lineages or in crosses between them. In Germany, we likely only have the western lineage sensu Bardy & al. (2010) but there may also be other genetic lineages present.

Albach & Greilhuber (2004) published the first genome size measurements for the species with 1C=0.896pg for diploids and 1C=1.487pg for tetraploids based on Feulgen densitometry. The latter value appeared to be too low based on the value for the diploids and subsequent values were higher (1C=1.86-1.93pg; Castro et al. 2012; Zonneveld 2019). Our values for the tetraploids are on average (1C=1.811pg) a bit lower but almost perfectly double the amount of the diploid value (Albach & Greilhuber 2004).

In preparation of a wider phylogeographical analysis, we have analyzed further populations but only found one diploid (Table 5). This is from a population in northern Hungary, close to a population sampled by Bardy et al. (2010). As populations from Central Germany appear to be tetraploid (Table 5), diploids are likely restricted to Southern Germany. Equally, measurements of populations originating from Poland and Lithuania revealed exclusively tetraploid plants. The only measurement from the Apennines reveals the presence of tetraploids in the region, even though the presence of diploids cannot be excluded, and it is plausible based on the pattern emerging in the species. Thus, diploids appear to be more prevalent at the southern margins of its distribution area, while tetraploids are spread throughout.

**Table 5:**
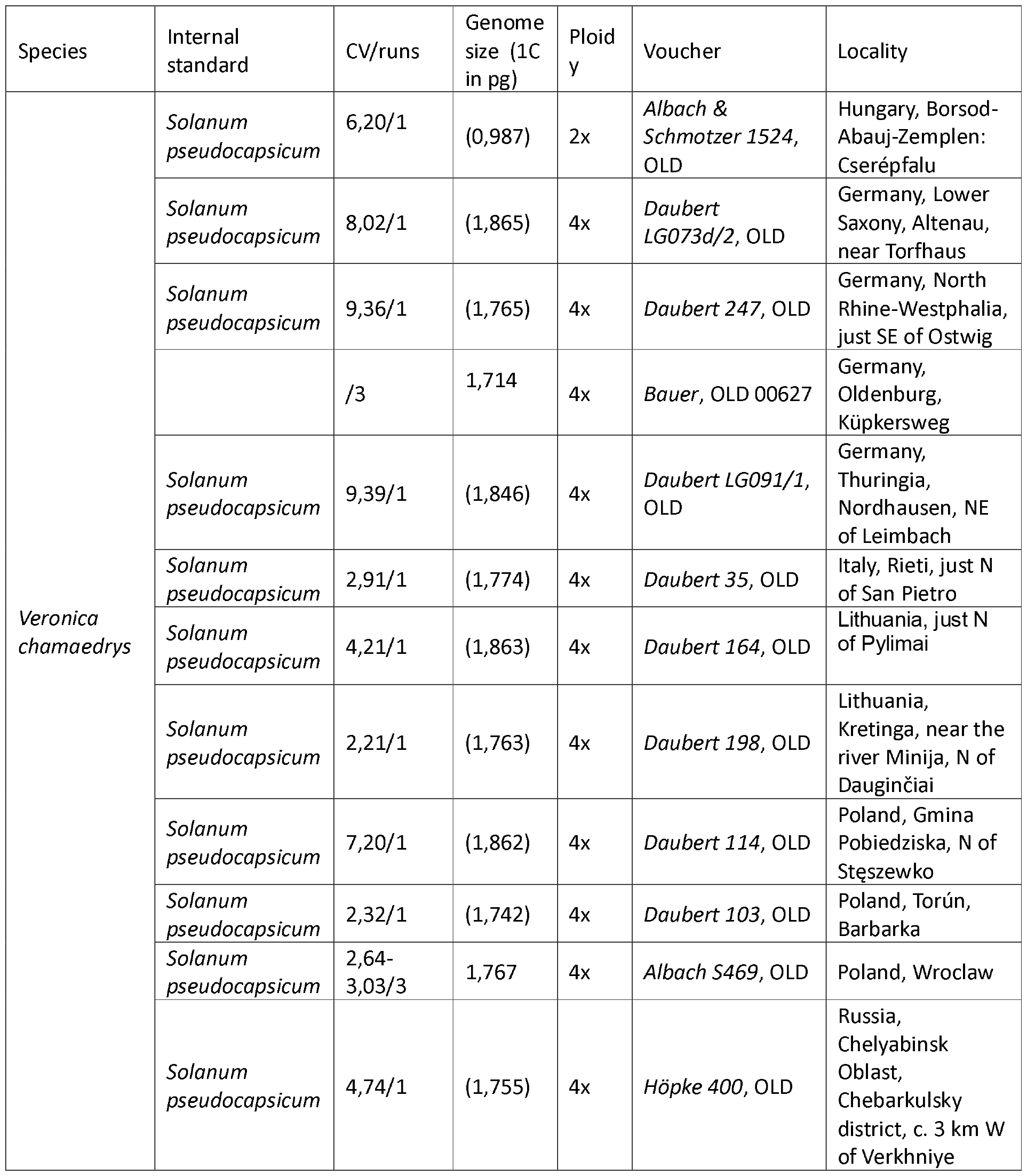
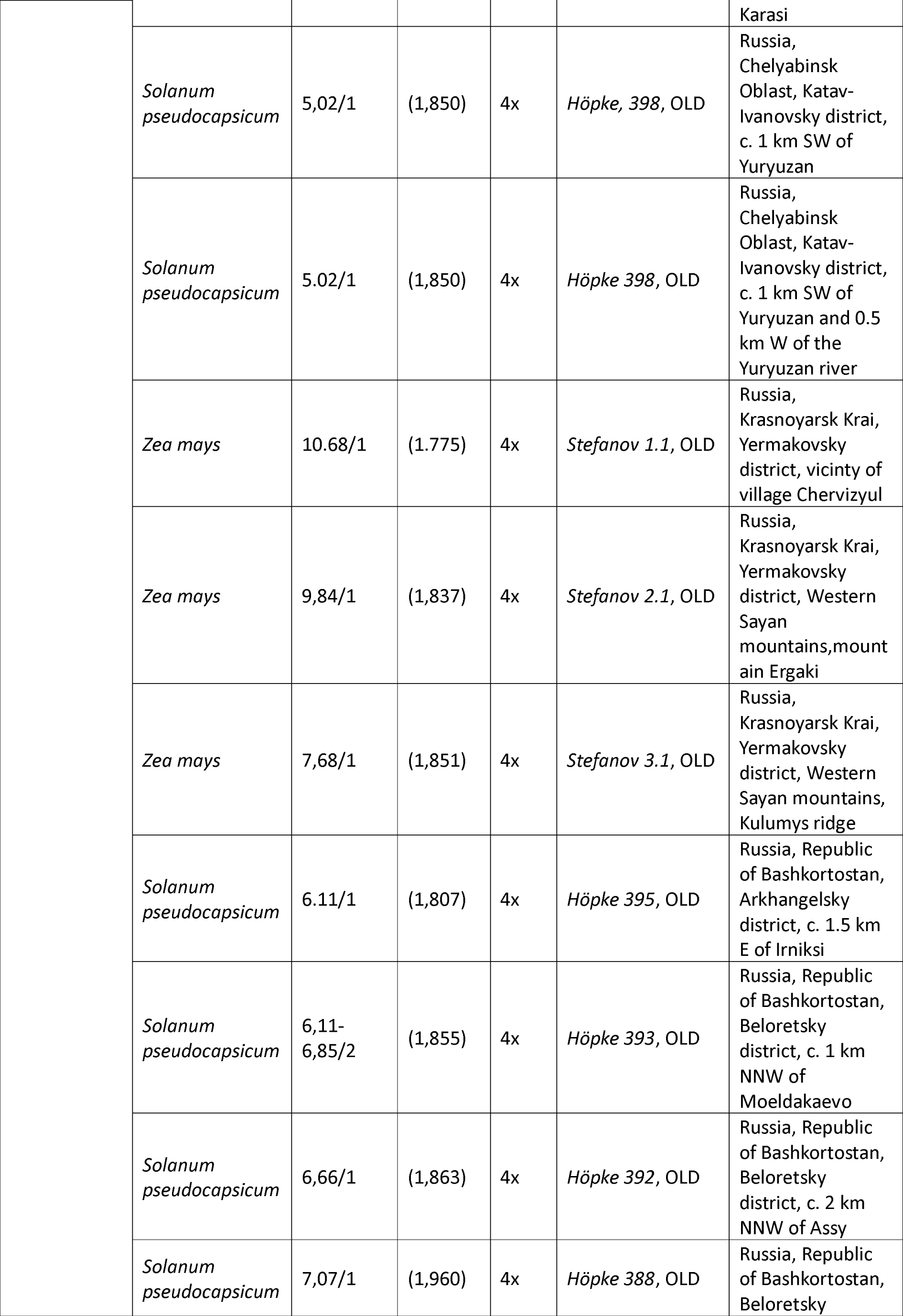
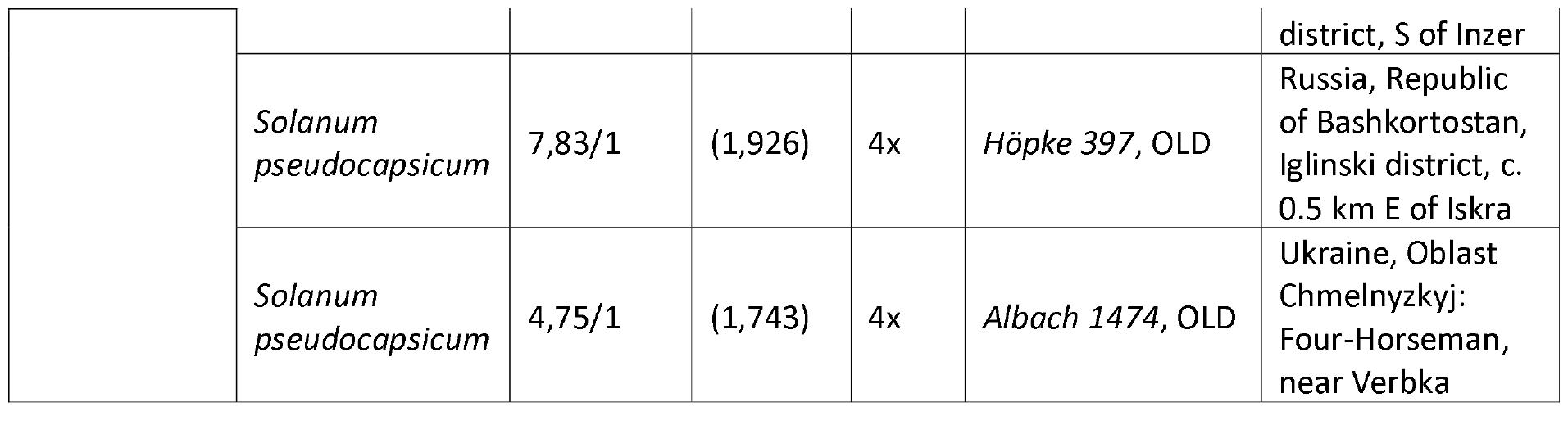
Information on genome size measurements for *V. chamaedrys*. Genome size measurements with only one run and/or cv above 7 were intended as tests of ploidy and should therefore not be considered reliable genome size estimates.

### 29. Veronica polita

*Veronica polita* is a widespread, weedy species that has been introduced in many countries of the world. Its chromosomes have commonly been counted and nine counts have been published later than Albach & al. (2008) adding up to 47 counts from Portugal to Japan with all but one being diploid with 2n = 14. Three counts are available from Germany with Tischler (1935) being of uncertain origin and Buttler & Buß in Gregor & Paule (2022) and Meve in Gregor & Paule (2023) confirming the diploid status of the species. Our genome size estimate is the fourth being published for the species, none from Germany. Our estimate (Table 6) and that of Castro & al. (2012) are somewhat lower than that of Albach & Greilhuber (2004) and Zonneveld (2019). The reason for this variation of almost 20% (1C= 0.36-0.43pg) is unclear.

**Table 6:**
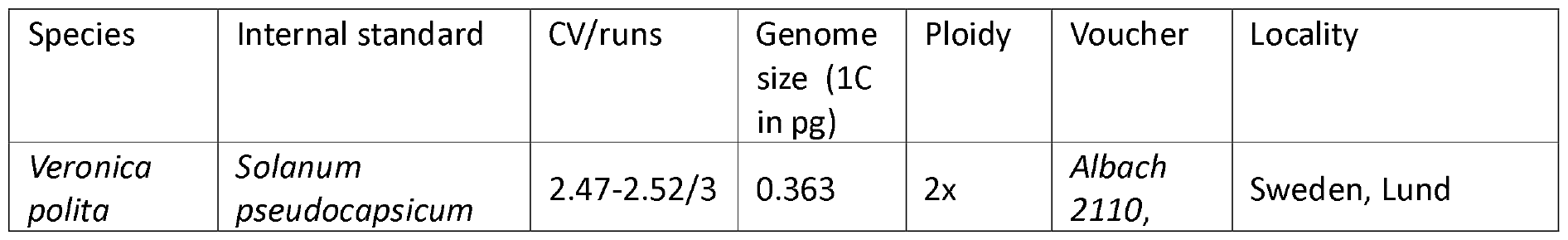
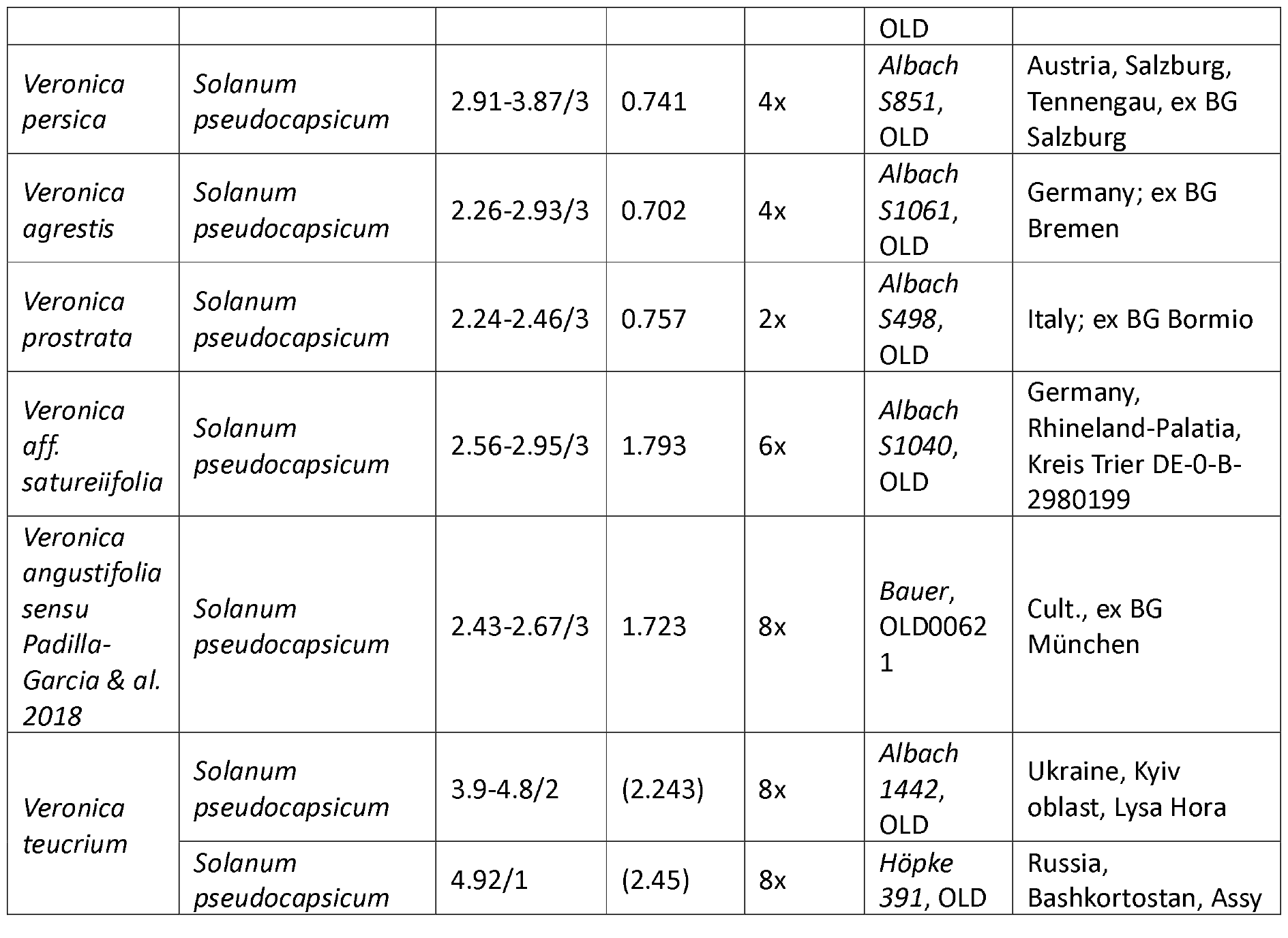
Information on genome size measurements for *V*. subgen. *Pocilla* and *Pentasepalae*. Genome size measurements with only one run and/or cv above 7 were intended as tests of ploidy and should be therefore not be considered reliable genome size estimates.

### 30. Veronica opaca

*Veronica opaca* is either an allopolyploid hybrid of *V. polita* and *V. bungei* (Fischer 1987) or an autotetraploid derivative of *V. polita* (Albach unpubl.). Consequently, all 12 counts so far show 2n = 28 chromosomes (Albach & al. 2008; Gregor & Hand 2009; Buttler in Paule & al. 2017). Unfortunately, no genome size measurement is available so far. Based on comparison with related tetraploid species, *V. persica* and *V. agrestis* (see below), a value of 1C=ca. 0.70-0.75pg is expected but if it is indeed an autotetraploid derivative of *V*. polita, the higher values of *V. polita* are correct, and it is indeed relatively young (Beatus 1936) with less time for genome downsizing, *V. opaca* may even have a somewhat higher genome size of more than 1C=0.8pg.

### 31. Veronica persica

The most common and widespread species in the genus is *V. persica*, a cosmopolitan weed, which explains why there are so many reports of its chromosome and throughout the world (Albach & al. 2008). Based on the uniformally counted 2n = 28 (Albach & al. 2008), it is tetraploid, and Fischer (1987) suggested that it is an allopolyploid cross of *V. polita* and *V. ceratocarpa*. The only count in Germany is from Tischer (1935). The previous four publications of genome size for the species, including one from the Botanical Garden Oldenburg, range between 1C=0.70-0.785pg (Bennett & Smith 1976, Castro & al. 2012, Zonneveld 2019, Meudt & al. 2015), with our new estimate being intermediate (Table 6). This is 12% intraspecific variation and tends to be a bit less than double the genome size of *V. polita*, which suggests that either *V. ceratocarpa* has a lower genome size than *V. polita* or genome downsizing has occurred.

### 32. Veronica agrestis

The third tetraploid species of *V*. subgen. *Pocilla* occurring in Germany is *V. agrestis*. 19 counts from Spain to Byelorussia supported 2n = 28 (Albach & al. 2008). Also, two counts from Germany (Wulff 1937, Buttler in Paule & al. 2017) support this number. Given the problems of Feulgen densitometry the first genome size measurement by Albach & Greilhuber (2004; 1C= 0.733pg) may have been too high (Meudt & al. 2015) and subsequent measurements are all lower, between 1C=0.69-0.71pg with those from Meudt & al. (2015) and ours here (Table 6) being from Northern Germany and the third from the Netherlands (Zonneveld 2019) while the initial count was from Greece (Albach & Greilhuber 2004).

### 33. Veronica filiformis

*Veronica filiformis* is a subalpine species of the Caucasus that became invasive in Europe and North America (Thaler 1953). Only one chromosome count exists from the native area (Aryavand 1975), whereas all other 12 counts are from the introduced area (Albach & al. 2008). All counts but none from Germany support a diploid status with 2n = 14 chromosomes. Albach & Greilhuber (2004) reported the genome size based on German plants as 1C= 0.37pg with the subsequent report by Zonneveld (2019) being a bit larger (1C=0.39pg). However, both measurements are in the same range as the other diploid species from *V*. subsect. *Agrestes, V. polita* (see above).

### 34. Veronica prostrata

*Veronica prostrata* s.str. is considered a purely diploid plant (Rojas-Andrés & al. 2015, 2020) with a distribution in central to eastern Europe, reaching west to Germany in Saxony-Anhalt to Bavaria. Four counts exist from Germany, all from Saxony-Anhalt (Frank & Hand in Gregor & Hand 2012; Dersch in Paule & al. 2017) confirming counts from other regions (Albach & al. 2008; Delgado et al in Marhold 2018). Genome size has not been measured from German plants but ranges between 1C=(0.70-)0.75-0.80(-0.88)pg (Rojas-Andrés & al. 2015, 2020; Table 6).

### 35. Veronica satureiifolia

*Veronica satureiifolia* was formerly considered synonymous with *V. prostrata* or its own subspecies under that name (*V. prostrata* subsp. *scheereri*) differentiated by some morphological characters (Brandt 1961) but also geography with *V. satureiifolia* occurring in western Europe from Spain to the Netherlands, reaching eastwards into western Germany (Rojas-Andres & Martinez-Ortega 2016). The distribution pattern reminds of the distribution of genetic lineages in *V. spicata* (see above), suggesting survival in common glacial refugia. Similar to *V. spicata* there is also a ploidy difference with the higher ploidy level occurring in the west. Scheerer (1937), Brandt (1961) and Albach & al. (2008) reported tetraploid plants (2n = 32) from Baden-Württemberg. Rojas-Andrés & al. (2020) reported plants of *V. satureiifolia* with 1C=1.29-1.40pg from France and northern Spain.

The biggest question in the taxonomy of *Veronica* in Germany is the identity of plants of *V*. subsect. *Pentasepalae* in Rhineland-Palatia and Saarland. Hand (2003) summarized the history of these plants and their morphology. The presence of *V. satureiifolia* (=*V. prostrata* subsp. *scheereri*) in the region is undisputed based on morphological similarities with *V. satureiifolia* in other regions. However, some authors suggest that another taxon exists, recognized as *V. orsiniana* by Hand (2003), a species otherwise present only in Italy, southern France and northeastern Spain. Other possible taxa are *V. angustifolia* (=*V. teucrium* var. *angustifolia*) sensu Padilla-Garcia & al. (2018) as suggested by Albach (2021) and *V. teucrium* var. *vahlii* (=*V. teucrium* subsp. *vahlii*) from southwestern Switzerland as suggested by Hartl (1975), a taxon synonymized with *V. orsiniana* by Walters & Webb (1972) and *V. angustifolia* by Padilla-Garcia & al. (2018). Hand (2023) recently and convincingly explained that *V. angustifolia* is not an appropriate name for the taxon and suggested the name *V. bastardii*. However, this taxon has mostly been used for plants belonging to *V. satureiifolia*. Therefore, a detailed morphological, karyological and molecular analysis is necessary to understand which taxa are present in northern France and adjacent areas such Rhineland-Palatia and Saarland. According to Hand (2003) the plants are prostrate with pubescent sepals and capsules, leaves that are intermediate in size between *V. satureiifolia* and *V. teucrium*. Given that the chromosome number is identical to that of *V. satureiifolia* (Hand 2003), chromosome numbers are not able to distinguish between *V. satureiifolia* and German *V. orsiniana* sensu Hand (2003). Further, all other counts of *V. orsiniana* are diploid (Albach & al. 2008; Rojas-Andrés & al. 2015, Delgado & al. in Marhold 2018) suggesting that it is distinct from our German plants. *Veronica angustifolia* sensu Padilla-Garcia & al. (2018) is octoploid (Delgado & al. in Marhold 2018; Table 6), which also precludes that it is the same taxon as the respective German plants, although its presence in Germany is possible (see below). In order to clarify the issue, we acquired seeds from the Botanical Garden Berlin from the region of Trier distributed as *V. austriaca* subsp. *vahlii*. The genome size of 1C=1.79pg (Table 6) corresponds to the hexaploid level. This supports the hypothesis by Hand (2003) that intermediate populations between *V. orsiniana* sensu Hand (2003) and *V*. teucrium exist in the Moselle valley and provides a new hypothesis. It is possible that *V. teucrium* (8x) and *V. satureiifolia* (4x) hybridize in the area to form hexaploid plants and backcross to *V. satureiifolia* to form the tetraploid plants of Hand (2003) with intermediate leaf size, prostrate habit of *V. satureiifolia* and pubescent sepals as in *V. teucrium. Veronica* subsect. *Pentasepalae* in southwestern Germany remains an important taxon to study.

### 36. Veronica teucrium

*Veronica teucrium* is octoploid with 2n = 64 (Albach & al. 2008; Rojas-Andrés & al. 2015) including several counts from Germany (Scheerer 1937; Dersch in Gregor & Hand 2014). It is the species of the genus with the highest chromosome number in Germany. Given the clear distinction from other German species by chromosome number but difficulties in always distinguishing it morphologically from *V*. austriaca, genome size estimates are important to distinguish species. The lowest genome size estimate for *V. teucrium* is that published here for a plant from Ukraine and that published by Albach & Greilhuber (2004) with values of 2.24pg and 2.25pg but most other estimates are about 2,30pg (Albach & Kosachev, in prep.) up to the value of 2,60pg reported here from Hungary. The estimate from Germany (1C=2.41pg) is intermediate (Rojas-Andrés & al. 2020). Other octoploid species of the genus in Europe are *V. sennenii* from northern Spain and *V. angustifolia* from western Europe (central and eastern France, western Switzerland and northern Italy). The genome size estimate published here (2,20 pg; Table 6) is at the lower end of those values seen in *V. teucrium* but also those of *V. angustifolia* reported by Rojas-Andres & al. (2020; 1C=2.35-2.75pg).

### 37. Veronica austriaca

Finally, *Veronica austriaca* is a hexaploid species (2n = 48) with an eastern to southeastern distribution area reaching west to Bavaria. Its occurrence in Baden-Württemberg is disputed (Rojas-Andrés & Martinez-Ortega 2016). Three subspecies are recognized but only subsp. *dentata* occurring in Germany (Rojas-Andrés & Martinez-Ortega 2016). Exceptions are tetraploids reported from Bosnia-Hercegovina (Delgado & al. 2018) and octoploids from Bulgaria (Peev 1978). Octoploids from Switzerland and Germany (Brandt 1952, 1961) should be, similar to the specimens from Baden-Württemberg mentioned by Rojas-Andrés & Martinez-Ortega (2016), more carefully investigated for confusion with *V. angustifolia* sensu Padilla-Garcia & al. (2018). No genome size estimate of *V. austriaca* from Germany is available but the estimates from other regions range between 1C=1.65-2.10pg without a difference between subspecies.

## Conclusion

Albach et al. (2008) reported that chromosome numbers are available for 310 of 448 species of *Veronica*. Since then, chromosome numbers have become available for an additional 10 species (Shiuchi & Kanemonot 2001; Singh et al. in Marhold & Kucera 2016; Delgado et al. in Marhold & Kucera 2018; Doostmohammadi et al. 2021; Ha et al. 2022) with another 10 species now newly recognized resulting in chromosome numbers now available for 320 of 458 species (70%). Chromosome numbers are available for all species present in Germany but only 22 (59%) are actually from German plants. For genome size measurements, which started to be gathered for *Veronica* by Bennett & Smith (1976), the situation looks a bit bleaker with values present for only 162 species (35%) worldwide (Albach, unpubl.). For species occurring in Germany estimates are available for all but two (95%) but estimates for plants from Germany are just 10 of 37 species (27%). For three species, intraspecific variation in ploidy in Germany is known (*V. chamaedrys, V. spicata, V. longifolia*) but for others it is possible (*V. cymbalaria*). Furthermore, for several species groups with subtle morphological distinction, ploidy information is an important character to verify species identification. We, here, provide the reference for such studies aiming to detect the differentiation of species complexes on the microscale or up to the regional scale. Study systems worth to pursue such studies are the microscale differentiation between *V. hederifolia* and *V. sublobata*, the regional distribution of taxa of *V*. subsect. *Pentasepalae* in southwestern Germany or the extension of diploid *V. chamaedrys* towards the north and west in Bavaria and possibly beyond.

## Acknowledgement

We are grateful to Silvia Kempen, Sabrina Schöngart and Kai Sohn for support in the lab. Part of this study was funded by the VW-Foundation (grant number 97771).

